# Optimizing SARS-CoV-2 RBD Boundaries for Enhanced *E. coli* Expression

**DOI:** 10.1101/2025.01.16.633331

**Authors:** Anamika Biswas, Arighna Sarkar, Sreejith Raran-Kurussi, Kalyaneswar Mandal

## Abstract

The outbreak of Coronavirus Disease 2019 (COVID-19) has posed a significant risk to global health, warranting the formulation of efficient preventive and therapeutic measures to tackle its causative agent, severe acute respiratory syndrome coronavirus 2 (SARS-CoV-2). The spike (S) protein of coronaviruses plays a pivotal role in viral attachment and entry into host cells. The receptor-binding domain (RBD) of the SARS-CoV-2 S protein has demonstrated a robust binding affinity to ACE2 receptors in humans. Consequently, it has become a prime target for therapeutic interventions using antibodies, vaccines, or other designed inhibitors. This paper presents an optimized RBD sequence that can be efficiently expressed in *Escherichia coli* and refolded to yield a functional protein. Using optimized refolding procedures, we obtained 10-12 mg of active protein from a one-liter LB culture. The biological activity of the refolded RBD was confirmed by monitoring its interaction with the designed LCB1 miniprotein ligand by surface plasmon resonance, wherein they exhibited significant affinity levels as reflected by their dissociation constants (K_D_s < 10 nM). The resulting RBD could be an ideal target for designing potent COVID-19 antivirals.

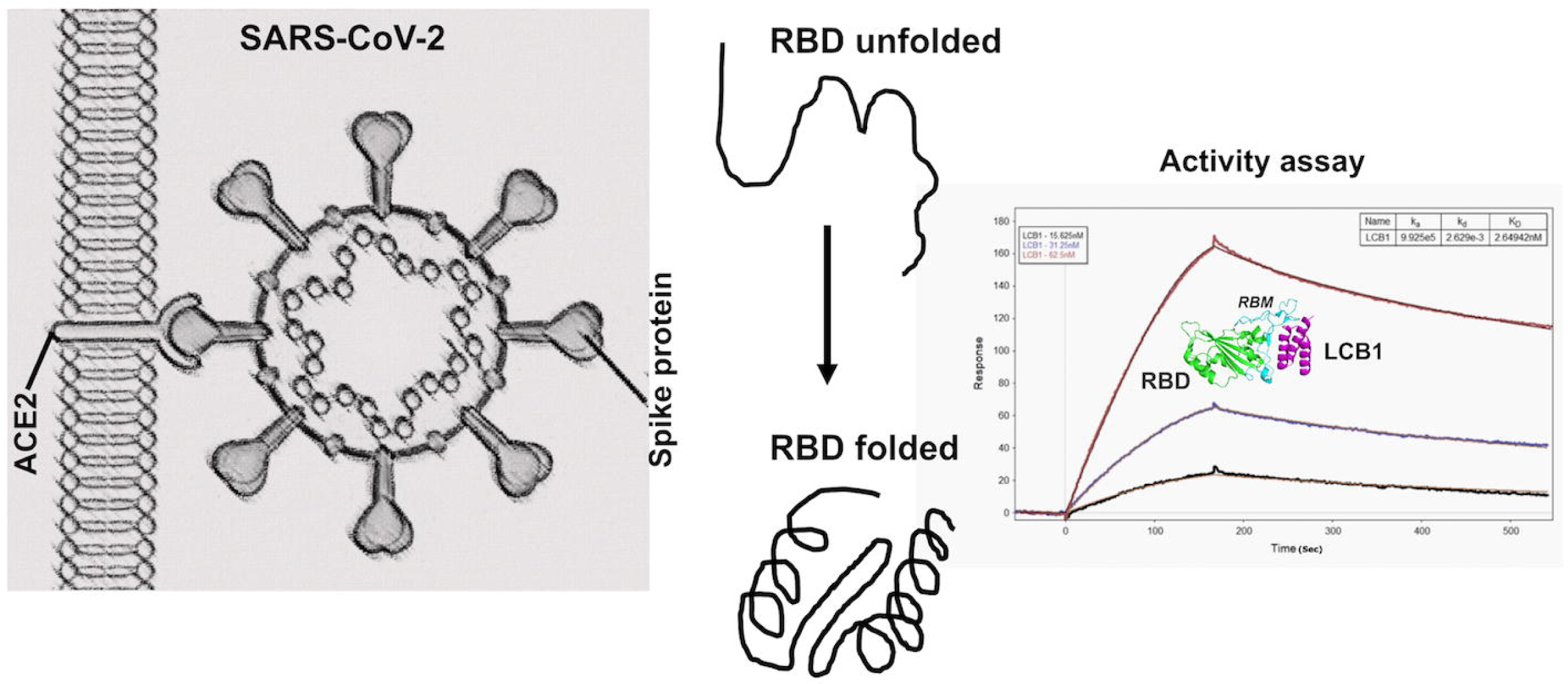

## 1. Introduction

Severe acute respiratory syndrome coronavirus 2 (SARS-CoV-2), a member of the SARS-related coronavirus family, is the etiological agent behind the COVID-19 pandemic, which has caused significant global disruption. The latest WHO report, as of November 2024, states a total of 777 million cases and approximately 7.1 million deaths (https://data.who.int/dashboards/covid19/deaths). The continuous evolution of SARS-CoV-2, driven by mutations, has led to numerous variants, posing a persistent challenge to public health (https://www.who.int/activities/tracking-SARS-CoV-2-variants). A robust therapeutic approach is essential to mitigate viral transmission and its associated impacts. Among the structural proteins of SARS-CoV-2, the spike (S) protein plays a pivotal role in viral entry into human cells. Its receptor-binding domain (RBD) specifically interacts with the angiotensin-converting enzyme 2 (ACE2) receptor, mediating infection [1–4]. This critical function makes the RBD a prime target for developing vaccines, neutralizing antibodies, and peptide-based therapeutics.

Efforts to express the RBD protein in both prokaryotic and eukaryotic systems have been extensively explored (Supplementary section, Table S1). While eukaryotic systems, such as insect and yeast cells, can produce soluble proteins, these often exhibit aberrant glycosylation patterns: O-glycosylation in insect cells and N-glycosylation in yeast cells, limiting their functional resemblance to the native protein [5]. These glycosylation inconsistencies, coupled with high production costs and low yields, restrict the utility of eukaryotic expression systems for applications such as serological testing.

Prokaryotic systems, particularly bacterial hosts, offer an alternative due to their cost-effectiveness and scalability. However, producing a functionally active and stable RBD protein in bacterial systems presents significant challenges, primarily associated with protein solubility and proper folding. Existing methodologies for bacterial expression and refolding of the RBD often yield suboptimal results, likely due to dependencies on sequence design and refolding conditions [6–13].

In this study, we address the challenges of producing functional RBD protein in bacterial systems by optimizing the RBD sequence for efficient expression and developing robust refolding protocols. By incorporating tags, such as maltose-binding protein (MBP) and/or 6X His, we achieved successful purification and refolding of the RBD, resulting in a biologically active protein with high functional efficacy. We present the optimized RBD sequence, describe the expression strategies employed, and provide a detailed refolding protocol. This work establishes a practical framework for producing functional RBD protein in bacterial systems, overcoming limitations of existing methods, and contributing to ongoing efforts in COVID-19 therapeutic development.

## 2. Materials and methods

### 2.1. Constructs

Based on the previously published reports, we have considered two sequence boundaries for RBD expression [2, 14]. The polypeptide chain from Arg^319^- Ser^591^ and Asn^331^- Ser^530^ is designated as RBD-1 and RBD-2, respectively (Figure 1). The pET-28a (+) expression plasmid with RBD-1 was custom-made (Genscript). The open reading frame (ORF) carried an N-terminal 6X His-tag and a TEV protease recognition site for tag removal. Additional gene cloning was done in-house to obtain the expression plasmids with different N and C-terminal tags.

**Figure 1.**
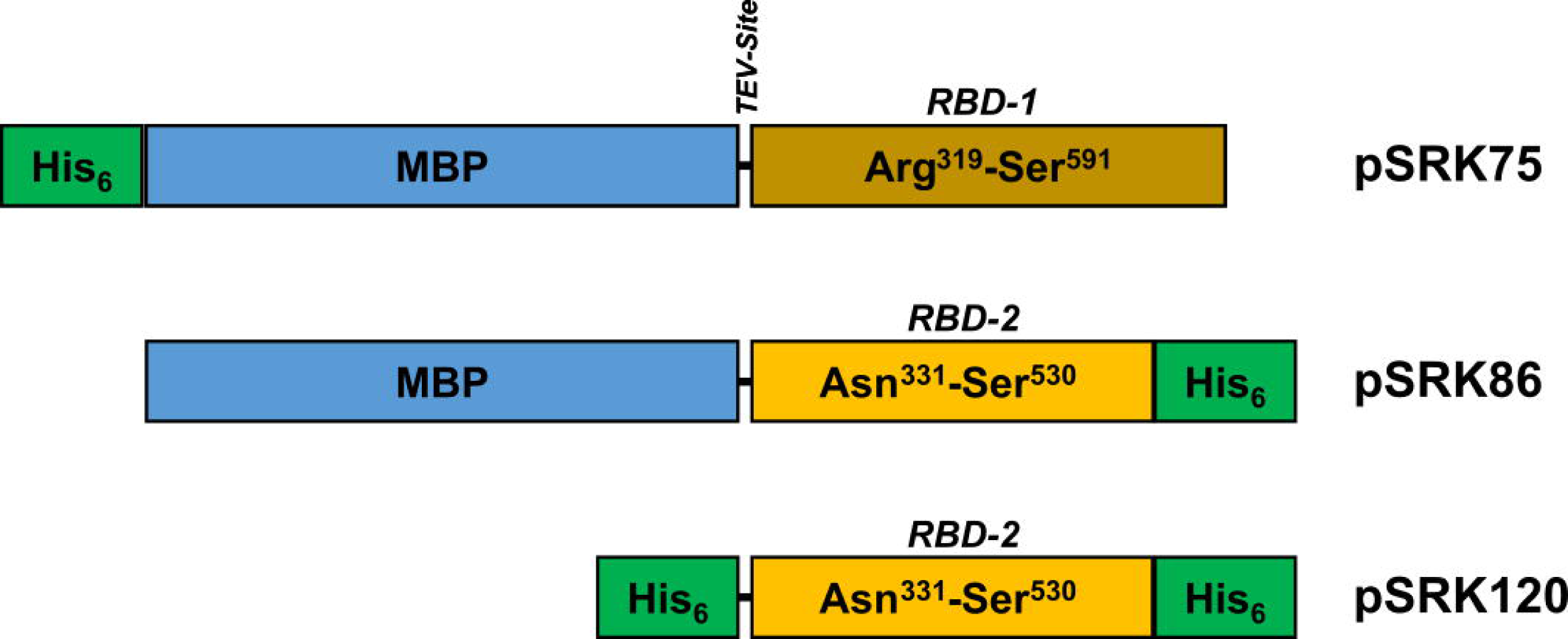
Schematic diagram of the expression vectors used in this study (not drawn to scale).

We used gateway cloning to generate the expression clones. The RBD ORFs encoding Arg^319^-Ser^591^ or Asn^331^-Ser^530^ sequence were amplified using appropriate primers (Table 1). The entry clones were made using a two-step PCR method. The ORF was amplified from pET-28a-RBD-1 in the first step using the RBD-specific forward and reverse primers. In the second step, the appropriate sequences for gateway cloning and TEV protease recognition were added using sequential PCR. The primers were designed to contain suitable overlaps for a seamless reaction. The final PCR product was recombined into pDONR221 to generate the entry clone. The standard BP and LR reactions were employed (ThermoFisher Scientific).

**Table 1:**
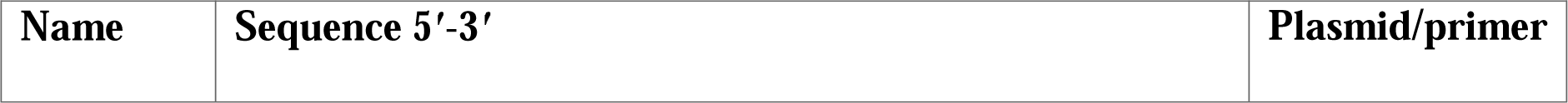
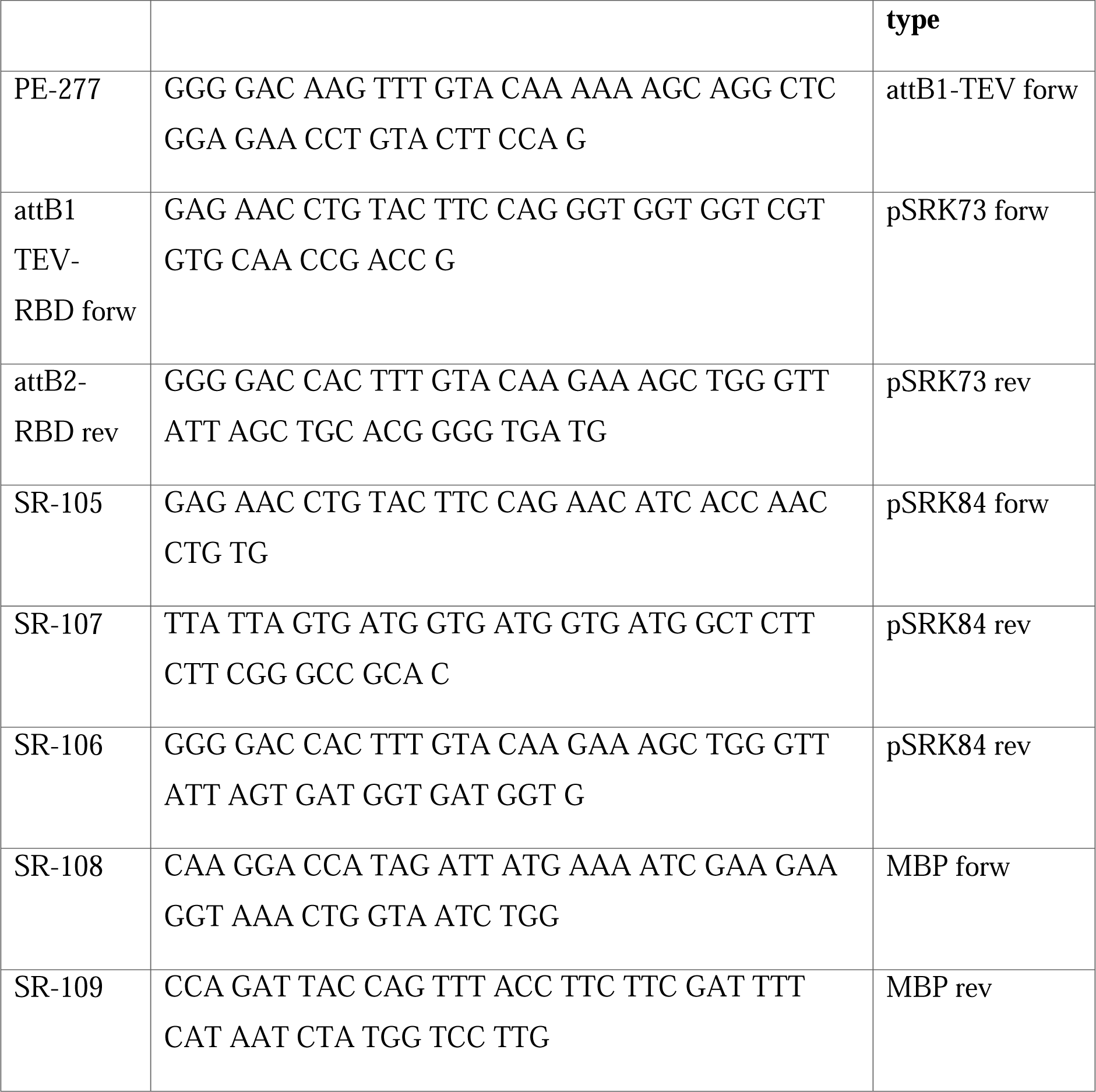
Primer sequences used in this study.

Specifically, entry clone pSRK73 (encodes Arg^319^-Ser^591^) was made using primers PE-277, attB1 TEV-RBD forw, attB2-RBD rev and was verified by sequencing. The primers are listed in Table 1. This design incorporates a TEV protease recognition site at the N-terminus of RBD. pSRK73 was subsequently recombined into pDEST-HisMBP to generate the expression vector pSRK75. Similarly, the pSRK84 entry clone (encodes Asn^331^-Ser^530^) was created using SR-105, SR-107, SR-106, and PE-277 primers using the pSRK73 template. This design incorporates a TEV protease recognition site at the N-terminus and a His_6_ tag at the C-terminus of RBD. The pSRK85 vector is a modified MBP destination vector lacking an N-terminal His_6_ tag. This construct was created using the pDEST-HisMBP template and the QuikChange Lightning Site-Directed Mutagenesis Kit (Agilent Technologies), with mutagenic primers SR-108 and SR-109. Subsequently, pSRK84 was recombined with pSRK85 and pDEST527 using an LR reaction to generate the expression vectors pSRK86 and pSRK120, respectively.

### 2.2. Protein expression

The expression of His_6_-tagged RBD was carried out in *E. coli* BL21 (DE3) cells or Rosetta 2(DE3), while MBP-tagged RBD constructs were tested in Lemo21(DE3) or SHuffle T7 Express cells. For soluble RBD protein production, bacterial cultures were grown in LB medium at 37 °C until the optical density at 600 nm (OD_600_) reached 0.7. Protein expression was induced with 1 mM IPTG, and the culture was incubated overnight at 18 °C, which improved protein solubility. For insoluble protein extraction, cultures were grown at 37 °C to OD_600_ 0.7, followed by IPTG induction. After 5 hours, cells were harvested by centrifugation, and inclusion bodies were isolated under denaturing conditions.

### 2.3. Purification of RBD from inclusion bodies

As the soluble proteins purified under native conditions were not functionally active (data not shown), proteins were extracted from the inclusion bodies under reduced conditions and refolded *in vitro*. We tested proteins made from His_6_-RBD-1 (GenScript custom-made), MBP-RBD-2-His_6_ (pSRK86), and His_6_-RBD-2-His_6_ (pSRK120) for *in vitro* refolding. The cell pellets were lysed in 6 M Gu.HCl + 2 mM BME + 20 mM Tris + 250 mM NaCl, pH 8, buffer. The homogenized cell suspension was sonicated, and the lysate was centrifuged for 1 hr at 15000 g. The supernatant was loaded onto a gravity column with Ni beads (Qiagen) for affinity purification. The column was washed with 5-10 column volumes (CVs) of wash buffer (8 M Urea + 2 mM BME + 20 mM Tris + 250 mM NaCl, pH 8,) and the bound protein was eluted using the elution buffer (250 mM imidazole + 8 M urea + 2 mM BME + 20 mM Tris + 250 mM NaCl, pH 8). The protein was obtained along with its BME adduct.

The mass of the eluted protein was confirmed by ESI-MS. After flash-freezing in liquid nitrogen, the protein was stored at -80 °C. However, during storage under these conditions, varying degrees of carbamylation were observed in the purified batches. Carbamylation typically results from the reaction of free amino groups (e.g., lysine residues or the N-terminal amino group) with isocyanic acid (HNCO), a byproduct of urea breakdown [15]. To mitigate this, we removed urea by dialyzing the protein against 10 mM phosphate buffer (pH 3) at 4 °C, with buffer changes at 2, 4, and 12 hours. This was done prior to lyophilization in subsequent preparations.

### 2.5. Refolding

For MBP-RBD-2-His_6_ (pSRK86-expressed protein), refolding by stepwise dialysis was carried out as described previously [16]. However, SPR analysis against LCB1 revealed that the refolded protein was inactive. For His_6_-RBD-2-His_6_ (pSRK120-expressed protein), folded via the stepwise dialysis method, we followed the same protocol mentioned previously with minor modifications (Supplementary section 3) [17].

After multiple trials, we also optimized a rapid dilution protocol for RBD refolding, which was successfully applied to both pSRK86 and pSRK120-expressed RBD. The final protocol is as follows. The lyophilized protein was dissolved in buffer (8 M Urea, 20 mM Tris, 100 mM NaCl, pH=8.4, concentration=5 mg/mL) and 10 mM DTT was added to it followed by incubation at 37 °C for 30 min to reduce the protein and eliminate BME adducts, if any. The protein was further dialyzed against 500 ml of 8 M urea + 20 mM Tris + 100 mM NaCl, pH 3 buffer for 12 h at 4 °C to reduce the imidazole concentration. To the dialyzed protein, EDTA was added to a final concentration of 1 mM. Next, this protein was added to the refolding buffer (20 mM phosphate + 0.5 M L-arginine + 2 mM GSH + 0.5 mM GSSG + 0.5 M urea + 100 mM NaCl, pH 8.0, 4 °C) in three separate portions, with a 12-hour interval between each addition. Before adding the protein into the refolding buffer for the first time, the buffer was flushed with nitrogen gas for about 15 min. After adding each portion, the nitrogen flush step was repeated for 5 min. The refolding buffer was stirred gently during the drop-wise addition of the protein, and the refolding mixture was left undisturbed at 4 °C. After adding the final portion of the reduced protein to the refolding buffer, the mixture was incubated undisturbed at 4 °C for 32-36 hours to optimize the refolding yield. The refolded protein was then concentrated to 5 mL using a stirred cell concentrator (Millipore) equipped with a 10 kDa cut-off membrane, and its mass was verified by ESI-MS. Prior to size exclusion chromatography (SEC) on a Superdex S-200 pg column (Cytiva) to remove misfolded species, the concentrated protein was dialyzed three times against 500 mL of 1X PBS to eliminate residual redox reagents. Following SEC, the concentration of each purified fraction was determined by measuring A280 using a NanoPhotometer (Implen). The molar extinction coefficient for each construct was derived from Expasy-ProtParam [18].

### 2.6. Chemical synthesis of LCB1

The LCB1 peptide (NH_2_-DKEWILQKIYEIMRLLDELGHAEASMRVSDLIYEFMKKGDERLLEEAERLLEEVER-αCONH2) [19] was synthesized on rink amide resin (substitution = 0.51 mmol/g) by solid phase peptide synthesis technique. Fmoc-protected amino acids were used to synthesize the peptide in a single stretch by an automated peptide synthesizer (Tribute UV-IR, Protein Technologies Inc). The protocol is detailed in the Supplementary section 2 (Figure S2).

### 2.7. Binding studies by SPR

Functional activity of the RBD protein was checked against its binding partner LCB1, using BI-4500AP SPR Instrument. The tagged protein was immobilized onto a Ni-NTA chip. Three channels were used for protein immobilization and one for reference. Binding assays were performed at 25 °C in pH 7.4 using 1X PBS + 0.005% tween-20 as a running buffer at a constant flow rate of 30 µl/min while immobilizing the protein on the chip. Before running a series of analyte concentrations over the flow cells at the same flow rate, 40 µM EDTA was added to the running buffer to minimize the non-specific interactions that may hamper RBD-LCB1 protein-protein interaction. The binding isotherm was obtained by allowing the analyte to flow over the immobilized protein for 150 sec. The dissociation curve was obtained by stopping the flow of the analyte while the running buffer continued to pass through the channels at the same flow rate. The final binding isotherm was attained by subtracting the reference channel and fitting the sensorgram using a 1:1 Langmuir adsorption binding isotherm. Average K_D_ was obtained from the three repeats of the binding experiments.

## 3. Results and discussion

### 3.1. Boundary optimization of RBD

The expression plasmid encoding the RBD-1 polypeptide, featuring a TEV cleavage site and a His_6_ tag at the N-terminus, was obtained commercially. The His_6_ tag at the N-terminus enables one-step affinity purification using immobilized metal affinity chromatography (IMAC) and facilitates binding assays via surface plasmon resonance (SPR), where the protein can be immobilized onto a Ni-NTA chip.

Our initial goal was to produce an active, soluble form of the protein. However, when bacterial expression failed to yield functional RBD-1 (His_6_-RBD-1 from Genscript), we developed a series of expression clones (pSRK75, pSRK86, pSRK120) by systematically choosing the tags and boundaries of the RBD open reading frame (ORF) (Figure 1). Each clone was designed based on insights gained from experimentally evaluating the preceding constructs. The first clone, pSRK75, carried an N-terminal fusion of His_6_-MBP to RBD-1. When pSRK75 failed to produce functional protein, we developed pSRK86 (MBP-RBD-2-His_6_), where a His_6_ tag was appended to the C-terminus of the MBP-RBD-2 to foster better binding to Ni^2+^, which successfully yielded the first batch of functional RBD protein. A key improvement involved trimming the RBD sequence at the C-terminal end to remove two cysteine residues, aligning it with the reported crystal structure to a major extent [14]. To verify if MBP is absolutely required for refolding, we further designed pSRK120 (His_6_-RBD-2-His_6_) devoid of MBP. We focused on optimizing the protocols using pSRK86 and pSRK120, as these were the only plasmids that yielded productive results. Data from unproductive plasmids were excluded from this study, ensuring that only the successful outcomes are presented.

### 3.2. Expression using pSRK86 plasmid

#### 3.2.1. Protein production under native conditions

A new sequence boundary, spanning Asn^331^ to Ser^530^ (RBD-2), was selected based on the reported crystal structure (PDB code: 6M0J, [14], which demonstrated that this region is sufficient for interaction with ACE2. MBP-RBD-2-His6 (pSRK86), featured an MBP tag at the N-terminus and a His_6_ tag at the C-terminus of RBD-2. At first, we attempted to express the fusion protein in soluble form in SHuffle T7 cells. We selected this strain, believing that these cells would be more capable of facilitating correct disulfide bond formation in the cytoplasm [20]. We faced several challenges, including poor binding to the nickel column and co-elution of a contaminating protein. Hence, we used an MBPTrap column (Cytiva) for the initial capture and followed it with an SEC run using a Superdex S-200 pg column (Supplementary Figure S1). The mass analysis of the purest concentrated fraction from this preparation revealed the presence of two proteins: a partially folded protein with a molecular mass of 65,614.61 Da and a bacterial protein, of 57 kDa (Figure 2). The 57 kDa protein band is most likely the GroEL chaperone (UniProt ID: P0A6F5; molecular weight = 57,197.66 Da). In our ESI-MS analysis, the observed molecular weight (57,199.46 Da) closely matched the calculated mass value of GroEL. It has been reported that MBP fusion proteins tend to associate with GroEL when they are partially or incorrectly folded during expression in *E. coli* [21]. The detection of GroEL in our preparation further reinforced this claim. Furthermore, surface plasmon resonance (SPR) analysis against LCB1 showed no detectable binding affinity, confirming that the soluble protein expressed from pSRK86 was misfolded and functionally inactive.

**Figure 2.**
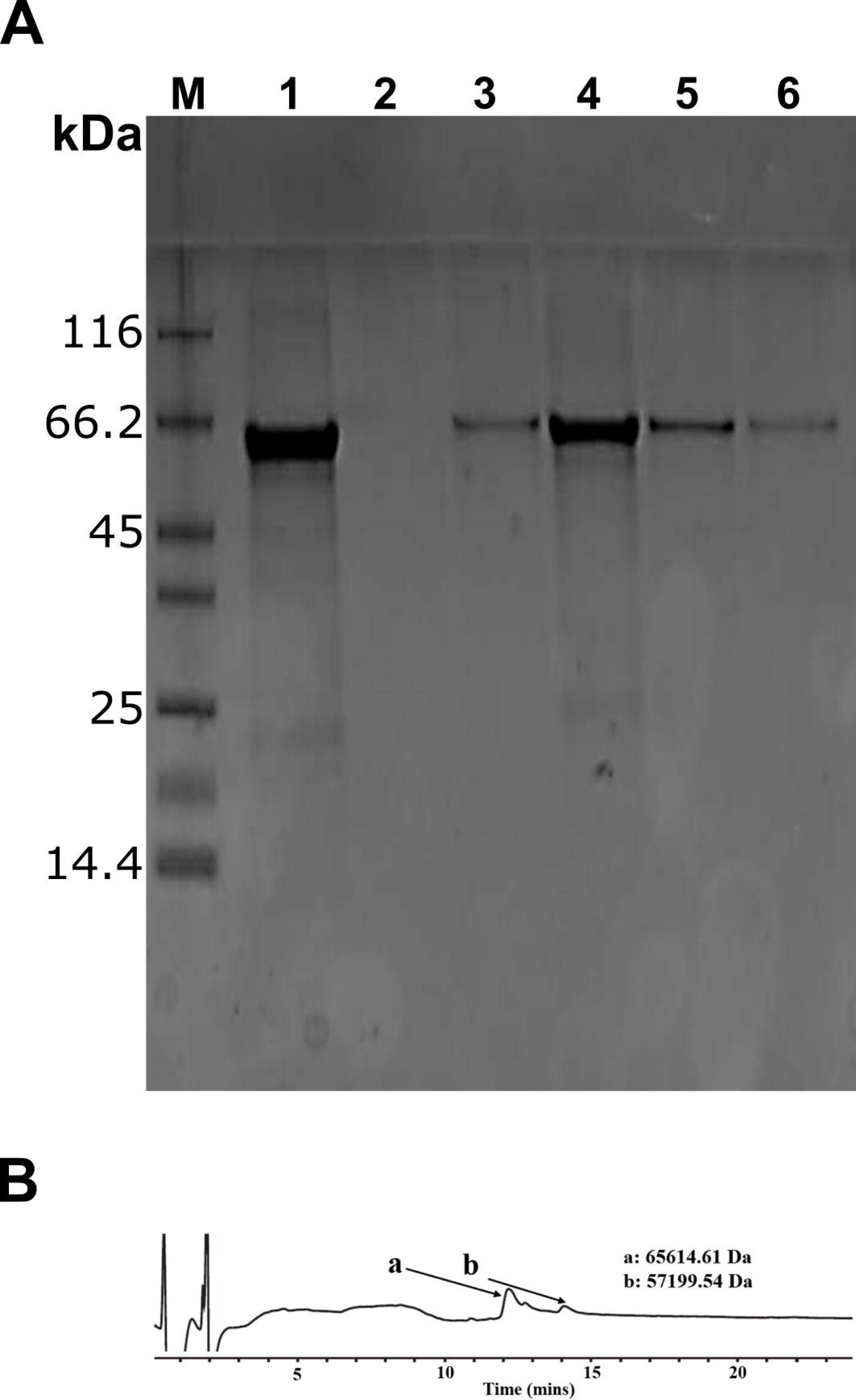
Soluble protein production using the pSRK86 plasmid. (A) SDS-PAGE analysis: Lane M, molecular weight marker; Lane 1, elution from the MBP-Trap column; Lanes 2–6, eluted fractions collected from the size-exclusion chromatography (SEC) column. (B) HPLC chromatogram of the purified soluble MBP-RBD-2 (the observed molecular masses are shown for the detected peaks).

#### 3.2.2. Functional protein production by rapid dilution refolding

Since purification under native conditions was unsuccessful, we employed *in vitro* refolding of MBP-RBD-2-His_6_ (pSRK86) using a rapid dilution method after isolating the protein from inclusion bodies (Figure 3). ESI-MS data confirmed the formation of four correct disulfide bonds after refolding. The authenticity of the proper protein fold was further validated by activity assays (see below). Typically, 5–10 mg was used in a refolding experiment, yielding 0.5–1.0 mg of active protein, corresponding to a 10% recovery. However, we note here that the refolding of the same protein by stepwise dialysis did not yield any active protein.

**Figure 3.**
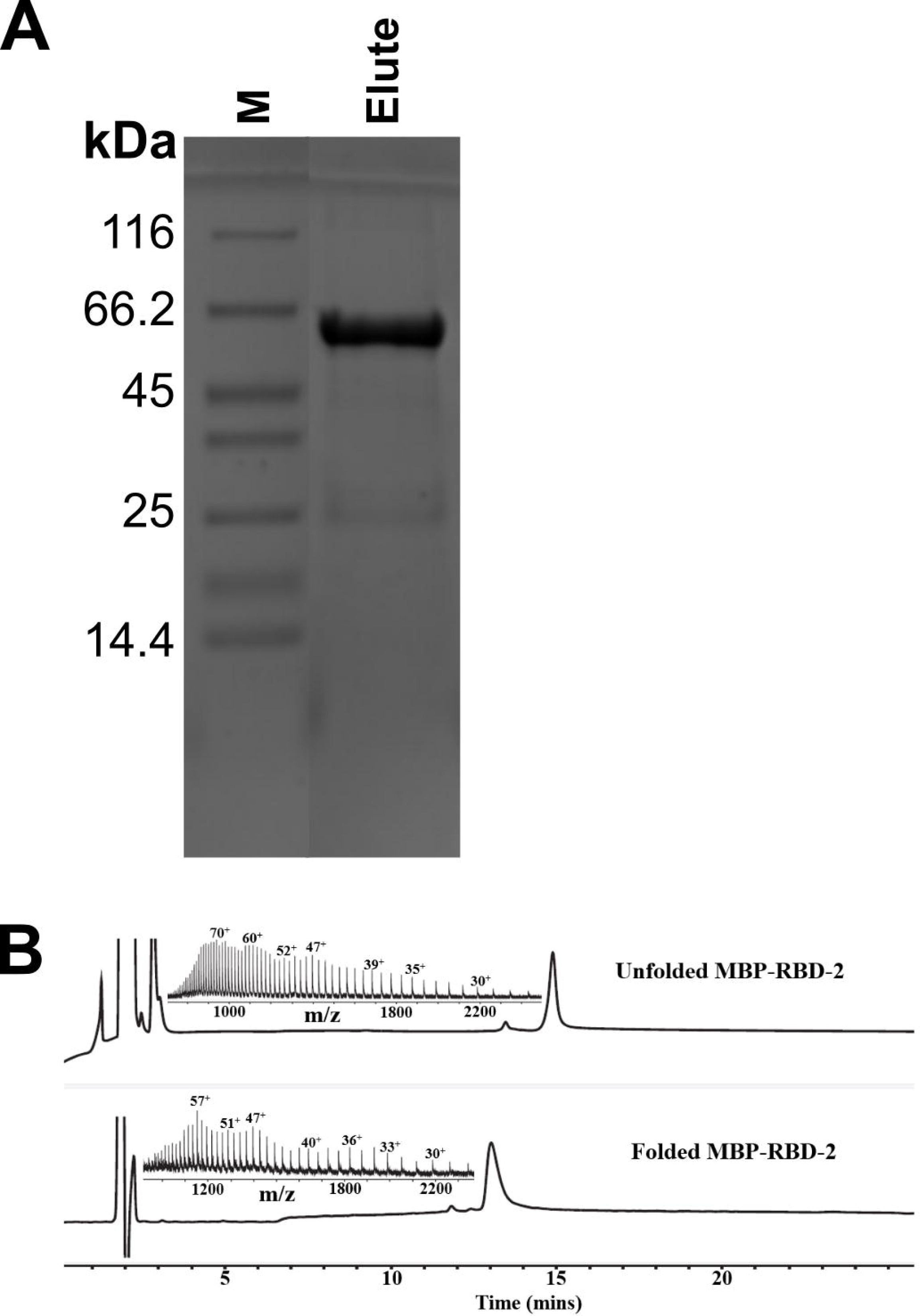
Purification of insoluble protein using the pSRK86 plasmid. (A) SDS-PAGE analysis: Lane M, molecular weight marker; Lane Elute, protein eluted from the HisTrap column under denaturing conditions. (B) ESI-MS data comparing the purified unfolded (65621.04 Da; top) and refolded RBD (65612.78 Da; bottom), alongside HPLC chromatograms.

### 3.3. Expression using pSRK120 plasmid

#### 3.3.1 Functional protein production by refolding

Refolding using rapid dilution and stepwise dialysis yielded comparable results for the His_6_-RBD-2-His_6_ from pSRK120 plasmid (Figures 4A and 4B). From 1 liter of bacterial culture, we typically purified 100-120 mg of protein under denaturing conditions. Of this, 5-10 mg was used in a refolding experiment, resulting in a 10% yield of the active protein (0.5-1.0 mg). Considering the smaller molecular weight of the tags present in this construct, the yield is significant, unlike the pSRK86 protein product wherein MBP contributes to the major part of the fusion protein. Both refolding methods successfully produced the active protein for the RBD expressed from pSRK120. A comparison of the unfolded vs folded charge states is shown in Figure 4C.

**Figure 4.**
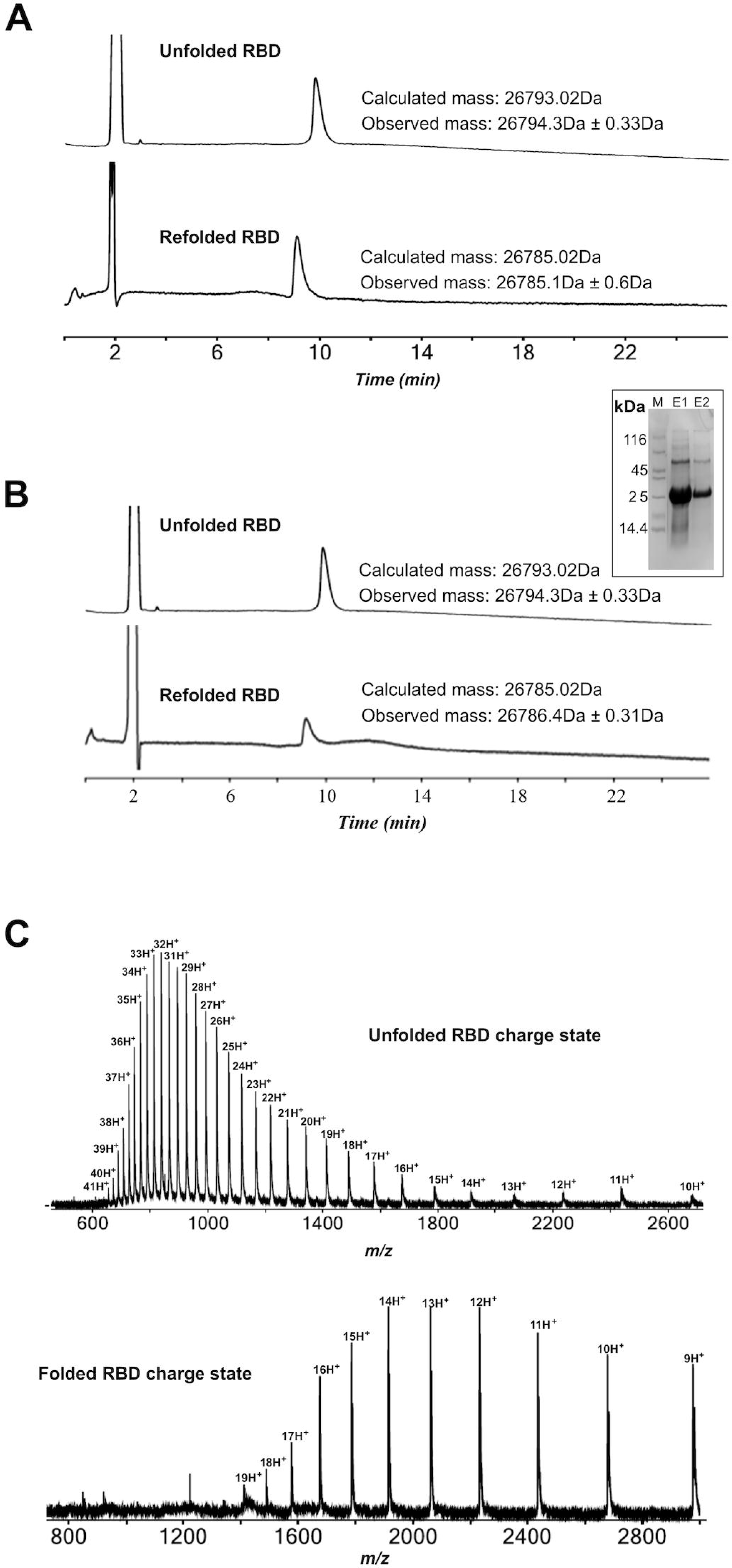
LC-MS analysis of RBD refolding using the pSRK120 plasmid. (A) HPLC chromatogram of the RBD refolded by the stepwise dialysis method. (B) HPLC chromatogram of the RBD refolded by the rapid dilution method. The calculated and observed masses are displayed in the inset. The purified protein (unfolded) analyzed by SDS-PAGE is also shown in the inset. Lane M, molecular weight marker; Lanes E1 and E2 carry protein eluted from the HisTrap column under denaturing conditions. (C) Mass spectrometry (m/z) data for unfolded and refolded RBD.

### 3.4 Activity of refolded proteins by SPR

Refolded proteins were immobilized on a Ni-NTA chip, and chemically synthesized LCB1 miniprotein, a strong binder of the SARS-CoV-2 RBD, was flowed over them at various concentrations. RBD from the pSRK86 plasmid showed minimal activity when refolded by stepwise dialysis (data not shown), but demonstrated strong LCB1 binding when refolded via rapid dilution (Figure 5A). The RBD expressed from pSRK86 bound to LCB1with a K_D_ value of 32.0 ±3.0 nM. ESI-MS data also confirmed the formation of the four disulfide bonds when we monitored the folding profiles before and after the refolding event. Interestingly, RBD expressed from the pSRK120 plasmid was active regardless of the refolding method used (Figures 5B and 5C). The protein showed K_D_ values of 3.02 ± 0.34 nM and 10.0 ± 1.0 nM for the refolded proteins from the stepwise dialysis and rapid dilution methods, respectively. The K_D_ data is presented as mean ± standard deviation (n = 3).

**Figure 5.**
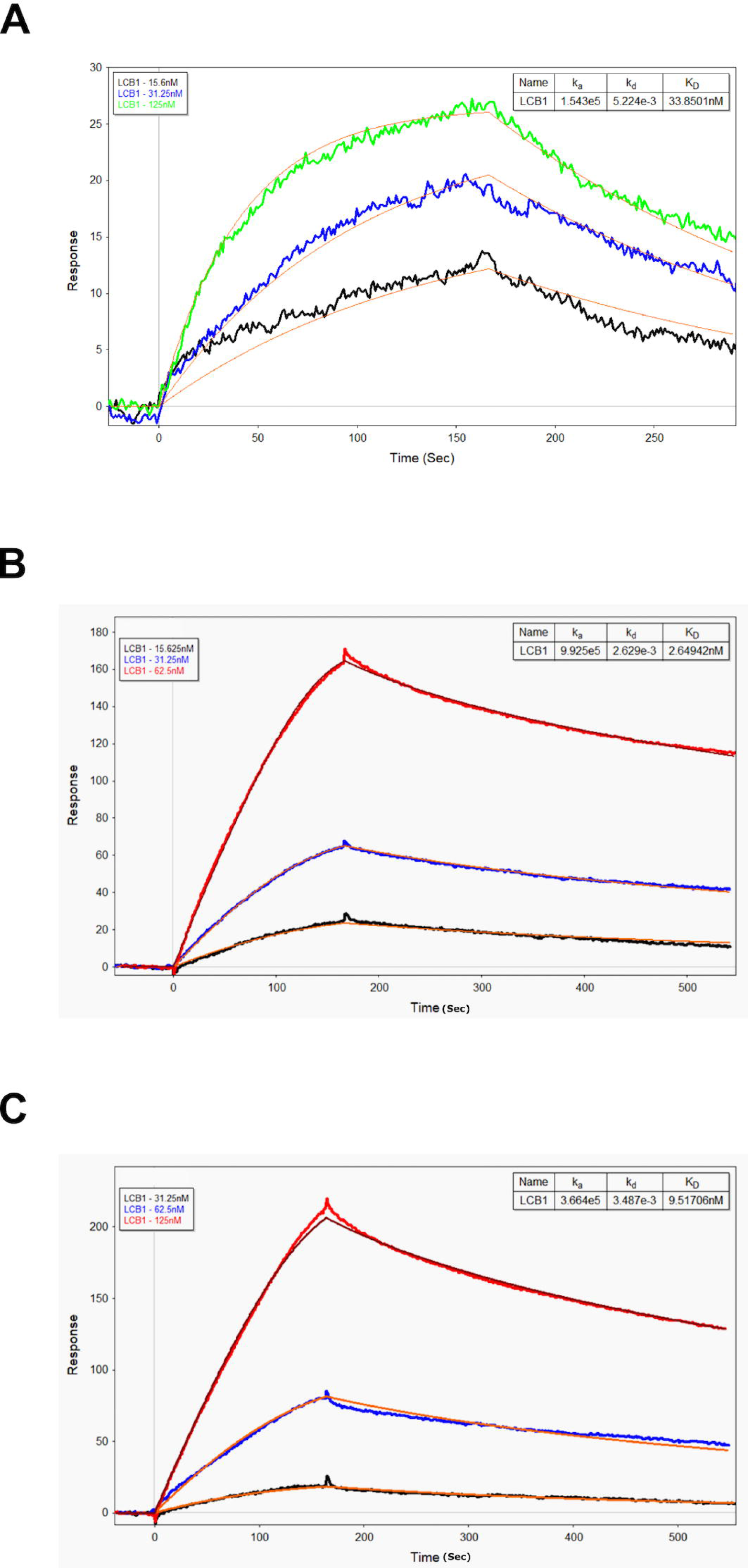
Activity Assay: SPR Kinetics of RBD Binding to LCB1. A) SPR sensorgram of the refolded protein (via rapid dilution) made using the pSRK86 plasmid. (B) and (C) show SPR sensorgrams of RBD refolded using the stepwise dialysis and rapid dilution methods, respectively, with protein expressed from the pSRK120 plasmid.

## 4. Conclusions

In summary, we successfully optimized the RBD sequence boundaries to enable efficient bacterial expression, achieving approximately 10% recovery of active protein after refolding. This was accomplished for both His_6_-tagged and MBP-His_6_ tagged fusion constructs. With large quantities of protein extracted from inclusion bodies, our primary focus was on the refolding process. From 1 liter of culture, 100-120 mg of denatured protein was obtained, yielding approximately 10-12 mg of active RBD after refolding. The His_6_-tagged version of the modified construct (pSRK120) showed enhanced activity, likely due to the absence of the hefty MBP tag, which may have hindered substrate binding in the pSRK86 variant. The crystal structure [14] suggests that the close proximity of the N- and C-termini in the pSRK120-expressed protein facilitates the effective display of the RBD binding surface by the His_6_ tags, thereby enhancing ligand binding in the SPR assay. Although N-terminal tag removal was technically feasible, we retained the tags for efficient immobilization for the SPR studies. Overall, our results indicate that the optimized sequence boundaries, rather than the tag, were crucial for achieving efficient refolding and high yields of active RBD. Nevertheless, we acknowledge that large protein tags may influence the refolding process, as seen with the RBD from the pSRK86 construct. While the fusion protein failed to refold effectively using stepwise dialysis, it was successfully refolded through rapid dilution, suggesting that the presence of the tags could play a role in the differing outcomes between the two protocols. The exact cause of this discrepancy remains unclear and will require further investigation through more rigorous experiments. Refolding a fusion protein necessitates more meticulous optimization than refolding the protein in its isolated form or with small peptide tags.

## Supporting information

Supplementary_section

## Acknowledgments

We gratefully acknowledge the support of the Department of Atomic Energy, Government of India, under Project Identification No. RTI 4007. The authors also thank the Biophysics Core Facility at TIFR Hyderabad for their valuable assistance.

