## Supplementary_section for "Optimizing SARS-CoV-2 RBD Boundaries for Enhanced *E. coli* Expression"

**1. Purification of native protein obtained from pSRK86 plasmid**

For native protein purification, the wet cell pellet (24.7 g) was resuspended in 250 ml of binding buffer (20 mM Tris + 200 mM NaCl + 1 mM EDTA, pH 7.52) with protease inhibitor cocktail tablets and 15 mg lysozyme. The cell suspension was stirred at 4 °C for 30 min and further sonicated with 4 short bursts of 2 min using the lowest power settings. Subsequently, the cell lysate was centrifuged (15000 g, 30 min, 4 °C) and the supernatant was filtered. Due to binding issues with Ni-column, we used an MBP-trap column (Cytiva) for initial capture since the soluble proteins carried an MBP tag. Further, the fusion protein was eluted with 20 mM Tris + 200 mM NaCl + 1 mM EDTA + 10 mM Maltose, pH 7.5 buffer. The pure fractions obtained were concentrated and loaded onto a Superdex S-200 pg size exclusion chromatography (SEC) column (Cytiva) for further polishing. We tracked the protein at all stages with SDS-PAGE (Figure S1).

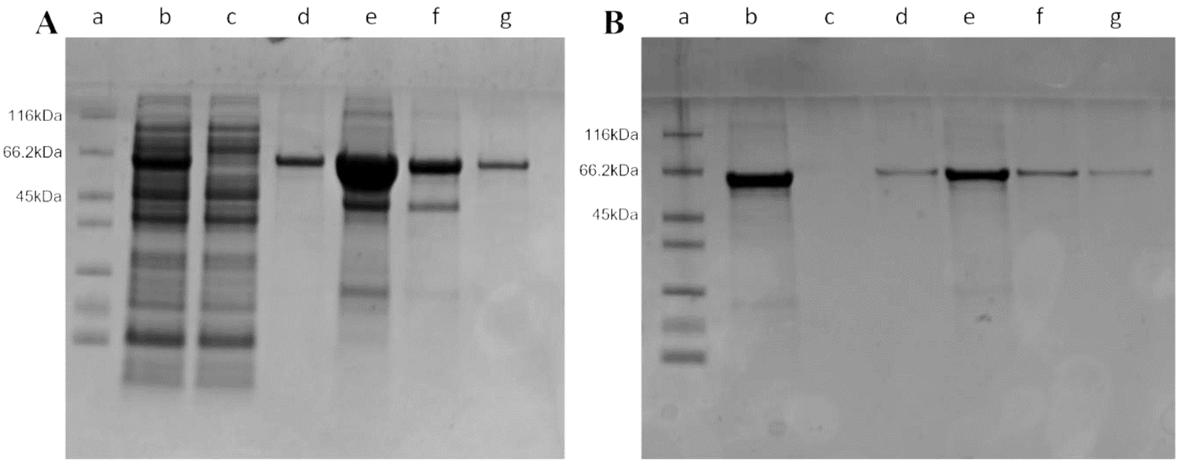


**Figure S1. SDS-PAGE analysis of native purification of the RBD expressed from pSRK86 plasmid. (**A) Purification of soluble protein using pSRK86 with MBP-Trap column. Lane a: molecular weight marker; b: load onto the column; c: flow-through; d-g: eluted protein fractions;

(B) Size-Exclusion Chromatography (SEC) profiles following MBP-Trap column purification. Lane a: marker; b: load onto Superdex-200 column; c-g: eluted protein fractions.

**2. LCB1 Miniprotein synthesis**

***2.1. Reagents***

N,N-Diisopropylethylamine (DIEA), N,N'-diisopropylcarbodiimide (DIC, HPLC grade) and all the $N^{\alpha}$ -Fmoc protected amino acids were obtained from Chem-Impex International. Dimethylformamide (DMF, HPLC grade), dichloromethane (DCM, HPLC grade), diethyl ether (AR grade), and trifluoroacetic acid (TFA, HPLC grade) were purchased from SRL Chemicals India. The HPLC grade and LCMS grade acetonitrile ($CH_3CN$) were purchased from Thermofisher Scientific. Piperidine was obtained from AVRA Chemicals, India. Fmoc-Rink-Amide resin was purchased from Supra Sciences, India. All other common reagents including triisopropylsilane (TIPS) and 3,6-dioxa-1,8-octane-dithiol (DODT) were purchased from Sigma-Aldrich and were of the purest grade.

***2.2. Reverse phase HPLC and LCMS analysis***

Analytical reverse-phase (RP) HPLC was performed on an Agilent HPLC instrument using an Agilent zorbax SB-C3 (5 µm), 4.6×150 mm reverse-phase silica column at a flow rate of 0.9 mL/min using a linear gradient of 10-70% solvent B in solvent A over 30 min at 40 °C (solvent A= 0.1% TFA in $H_2O$; solvent B = 0.08% TFA in acetonitrile). The UV absorbance of the column eluent was monitored at 214 nm wavelength. The peptide masses were measured over the entire UV absorption peaks corresponding to the compounds characterized by on-line LC-MS using an Agilent 1290 infinity II/6530 Q-TOF LC/MS instrument. The deconvolution of the charge states of the observed mass was carried out using Agilent MassHunter Qualitative Analysis software (version B.07.00). Calculated masses were based on average isotope composition or based on the most abundant isotopologue mass determined from the isotopic distribution provided by Agilent MassHunter Qualitative Analysis software.

Preparative reverse phase HPLC (RP-HPLC) of the crude peptide was performed with an Agilent ZORBAX-SB C3 (5 µm, 80 Å, 9.4 × 250 mm) column using an appropriate shallow gradient (15-45% solvent B in solvent A over 60 min at 40 °C; solvent A= 0.1% TFA in $H_2O$; solvent B = 0.08% TFA in acetonitrile) at a flow rate of 5 mL/min. Fractions containing the purified target peptide were identified by ESI-MS. The pure fractions were then pooled and lyophilized.

### *2.3. Protocol for machine-assisted Fmoc-SPPS and peptide cleavage from resin*

Peptide was synthesized using an automated peptide synthesizer (Tribute-UV/IR from Protein Technologies, USA). Fmoc-SPPS was carried out using amino acids (AA) (0.25 M), DIC (0.25 M) as a coupling reagent, and Oxyma (0.25 M) with DIEA (0.025 M) as additives. All amino acids except Arginine and Histidine were coupled for 10 min at 70 °C. Arginine was coupled for 20 min at room temperature followed by 5 min at 60 °C. Histidine was coupled for 5 min at room temperature followed by 12 min at 50 °C. Fmoc deprotection after every coupling cycle was carried out by 20% piperidine treatment at 40 °C for 1 min followed by 3 min at room temperature. After synthesis, the peptides were cleaved from the resin using TFA (85%), phenol (5%), TIPS (2.5%), water (2.5%), DODT (2.5%) and thioanisole (2.5%) as a cleavage cocktail. After cleavage, the TFA was evaporated under $N_2$ flow inside a well-ventilated fume hood. The cleaved peptide was precipitated by adding cold diethyl ether. The resulting precipitate was washed twice with cold diethyl ether and the crude peptide was lyophilized. The lyophilized powder was subsequently dissolved in water, pH adjusted to 2 using 6 N HCl, and loaded directly onto a preparative HPLC column for purification.

*2.4. Synthesis of LCB1 miniprotein*

The LCB1 miniprotein having the sequence as follows,

$NH_2$-DKEWILQKIYEIMRLLDELGHAEASMRVSDLIYEFMKKGDERLLEEAERLLEEVER-$^{\alpha}$CONH2 was synthesized on Rink-Amide resin (substitution = 0.51 mmol/g) by stepwise Fmoc chemistry SPPS on a 0.2 mmol scale in an automated peptide synthesizer. Purification using preparative HPLC afforded the pure peptide. Observed mass (ESI-MS): 6809.54 ± 0.03Da (average deconvoluted mass); calculated mass (average isotope): 6809.48 Da.

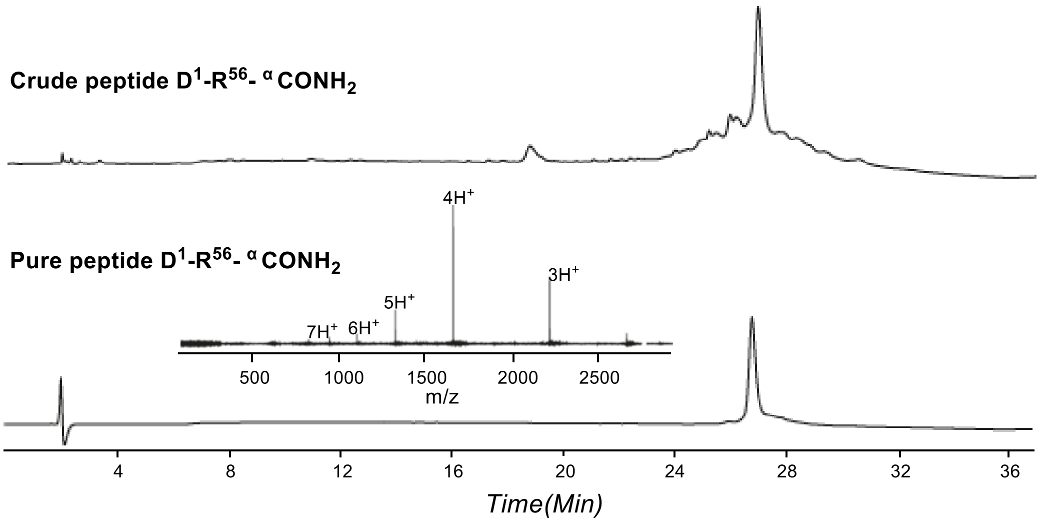


**Figure S2. LC-MS chromatogram profile and Mass spectrometry (m/z) data of synthesized LCB1 miniprotein.**

**3. Stepwise dialysis refolding of pSRK120 encoded protein**

The RBD protein ($His_6$-RBD-2-$His_6$) obtained after Ni-column purification from inclusion bodies was consecutively dialyzed against approximately 1.5 L of 10 mM $NaH_2PO4$ (pH 3.0, 4 °C) for 2, 4, and 12 hours. The precipitated protein was collected by centrifugation, lyophilized, and stored at -20 °C for future use. For refolding, approximately 5 mg of lyophilized protein was dissolved in Buffer B (6 M Gu.HCl, 50 mM Tris.HCl, 10 mM DTT, pH 8.0) to get a final concentration of 0.5 mg/mL and incubated at 37 °C for 30 min to reduce BME adducts. The solution was then dialyzed against 500 mL of Buffer C (6 M Gu.HCl, 50 mM Tris.HCl, pH 8.0) at 4 °C for 12 hours to remove excess DTT and BME. This was followed by a series of dialysis steps against 500 mL of various buffers at 4 °C as follows. Buffer D (6 M Gu.HCl, 50 mM Tris.HCl, 3 mM GSH, 0.9 mM GSSG, pH 8.0) for 24 hours, Buffer E (2 M Gu.HCl, 50 mM Tris.HCl, 400 mM L-arginine, 3 mM GSH, 0.9 mM GSSG, pH 8.0) for 36 hours, and Buffer F (1 M Gu.HCl, 50 mM Tris.HCl, 200 mM L-arginine, 3 mM GSH, 0.9 mM GSSG, pH 8.0) for 36 hours. The protein was then dialyzed against 500 mL of Buffer G (50 mM Tris.HCl, 100 mM L-arginine, 250 mM NaCl, 3 mM GSH, 0.9 mM GSSG, pH 8.0) for 24 hours and finally against 500 mL of Buffer H (50 mM Tris.HCl, 150 mM NaCl, pH 8.0) for 24 hours at 4 °C. After the final dialysis, the solution was centrifuged to remove precipitates, and the supernatant was concentrated to 5 mL using an Amicon 10 kDa membrane. The concentrated protein was purified by size-exclusion chromatography on a Superdex 200 pg column (Cytiva) to remove soluble misfolded aggregates. The concentration of pure fractions was determined by A280 measurement (Extinction coefficient, E=36830 $M^{-1}cm^{-1}$) using a NanoPhotometer (Implen, Germany). The refolding process yielded approximately 10%, with 0.48 mg of properly folded protein recovered from 5 mg of starting material. For functional analysis, the refolded protein was immobilized on a Ni-NTA chip and assessed by SPR for binding to LCB1.

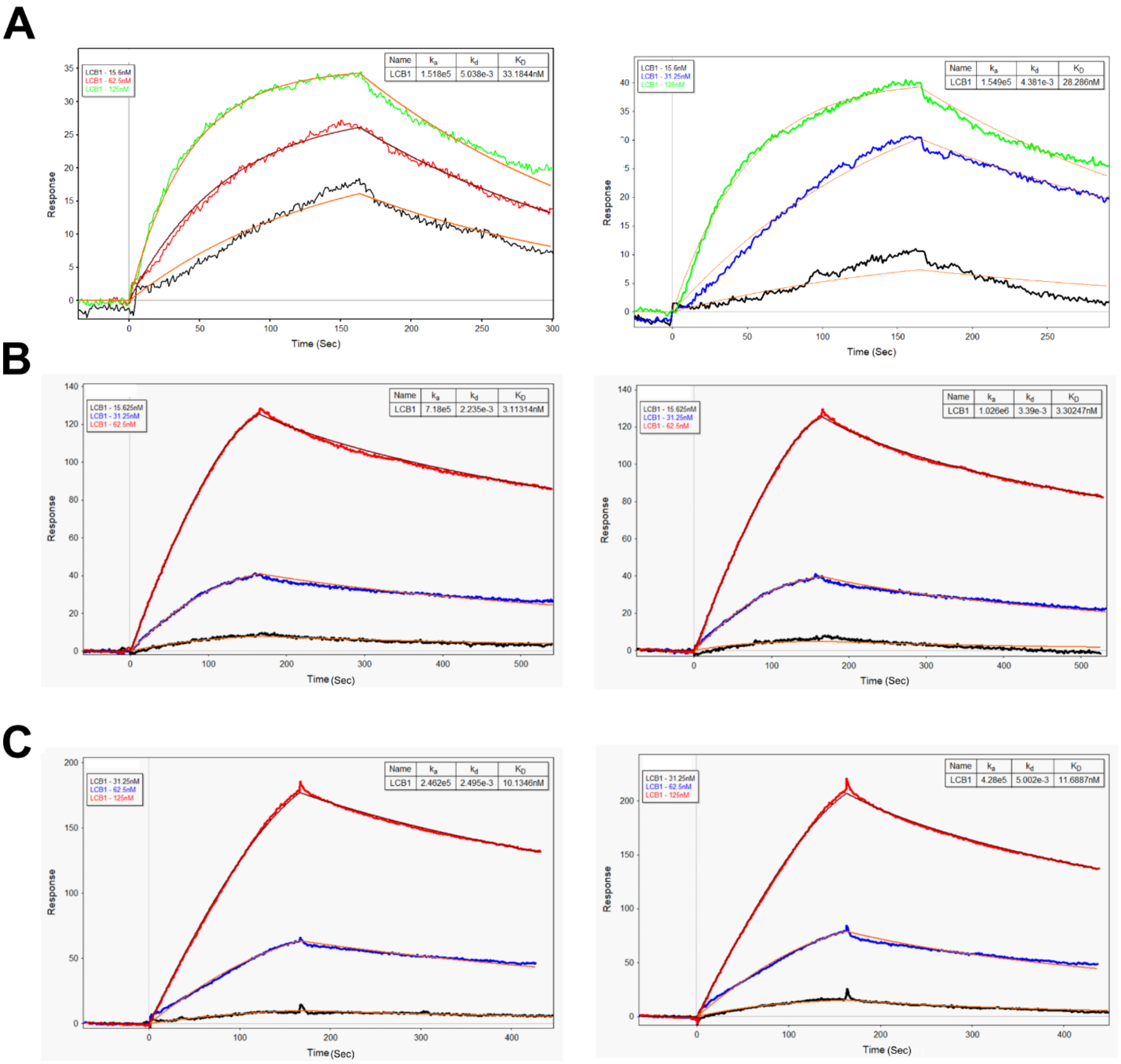


**Figure S3. Activity Assay: Kinetics of RBD Binding to LCB1 measured by SPR.** (A) SPR sensorgram showing the binding kinetics of RBD refolded via rapid dilution, expressed using the pSRK86 plasmid. (B) SPR sensorgram of RBD expressed from the pSRK120 plasmid and refolded using the stepwise dialysis method. (C) SPR sensorgram of RBD expressed from the pSRK120 plasmid and refolded using the rapid dilution method. These experiments replicate and extend the results presented in Figure 5 of the main text.

**Table S1. Summary of previous reports on RBD production**

| **Expression Host** | **Yields** | **Activity Assay details** | **Reference** |
|---|---|---|---|
| *Pichia pastoris* | 10-13 mg/L | Immunization studies | [1] |
| HEK-293T cells | 5 mg/L | Immunization studies | [1] |
| *E. coli* | Not mentioned | Binding to ACE2 on cell surface | [2] |
| *E. coli* | Not mentioned | SPR binding to ACE2 $K_D$=29.8 nM | [3] |
| *E. coli* | Not mentioned | ELISA assay | [4] |
| *E. coli* | 1.5 mg/L | ELISA assay | [5] |
| Green Algae *Chlamydomonas reinhardtii* | 1.8 ug/gram of wet biomass | ACE2 binding reporter assay $EC_{50}$=36 nM | [6] |
| *E. coli* | 0.25 mg/L | SPR against ACE2 $K_D$=19.01 nM | [7] |
| *E. coli* | Not mentioned | SPR against ACE2 $K_D$=10.4 nM | [8] |
| *E. coli* | 2 mg/L | Binding to ACE2 by QCM analysis $K_D$=180 nM | [9] |

| | | | |
|---|---|---|---|
| *E.coli* (Omicron variant) | Not mentioned | ELISA and BLI binding to ACE2 $K_D$=10.1 nM | [10] |
| *E. coli* | 90 mg/L | No binding observed with ACE2 | [11] |
| CHO-K1 cells | 50-100 mg/L | BLI binding to ACE2 $K_D$=58 nM | [11] |
| CHO-cells | 40 mg/L | ACE2 binding by Flow cytometry | [12] |
| *E. coli* | 2 mg/200 mL LB culture | BLI binding to ACE2 $K_D$=7.95 nM | [13] |
| *E. coli* (Beta, Delta, and Omicron variants) | 1 mg/L (Beta)<br>0.825 mg/L (Delta)<br>0.4 mg/L (Omicron) | ELISA assay for binding with ACE2 | [14] |
| *E. coli* (Wuhan, Alpha, Beta) | 0.5 mg/g of cell paste (Wuhan)<br>0.2 mg/g of cell paste (Alpha)<br>0.15 mg/g of cell paste (Beta) | BLI binding to ACE2 $K_D$=3.5 nM | [15] |
| *E. coli* (Omicron BA.5) | 3 mg/200 mL culture | BLI binding to ACE2 $K_D$=0.83 nM | [16] |

| | | | |
|---|---|---|---|
| *E. coli* (2 variants) | 1.8 mg/L or 4.5 mg/L | Binding to virus-neutralizing antibodies | [17] |
| *E. coli* | 6 mg/L | ELISA assays to antibodies and flow cytometry binding to ACE2 | [18] |
| *E. coli* (Omicron) | Not Mentioned | BLI binding to ACE2 $K_D$=0.83 nM | [19] |
